# Improving the classification of wildlife conservation status to support nature protection in the European Union

**DOI:** 10.1101/2025.07.01.662537

**Authors:** Marco Davoli, Martin Jung, Piero Visconti, Carlo Rondinini, Alessandra D’Alessio, Michela Pacifici

## Abstract

Ensuring that species of conservation concern achieve favorable conservation status (FCS) is central to European Union (EU) biodiversity conservation targets. A key criterion for FCS is exceeding the favorable reference range (FRR)—the range extent needed for long-term species stability and full ecological variation. However, due to data limitations, FRRs are often unknown, undermining their applicability. We developed a machine-learning approach to estimate and standardize FRRs across the EU. Applied to amphibians, mammals, and reptiles, our method provided FRRs for 99.5% of species of conservation concern, compared to 17.5% previously available (with satisfactory modelling performance: R^2^ 0.75). We reassessed conservation status using the estimated FRRs, finding that species in FCS (34.8%) are notably fewer than reported in official documentation (69.1%). The average proportional distance to FRR for species in unfavorable conservation status is -64.4%. Our approach may support periodic FCS reassessments and help refine the targets of EU conservation policies.

## 1 Introduction

The European Union (EU) has made notable strides in nature protection through continent-wide regulations such as the Habitats and Birds Directives, the Natura 2000 Network, and, more recently, the EU Biodiversity Strategy to 2030, which includes the Nature Restoration Regulation (Lang 2023). However, biodiversity continues to decline within the EU (European Environment Agency, 2020), primarily due to inadequate coordination among EU Member States, poor integration with other policies, insufficient funding, and limited stakeholder engagement (Hermoso et al. 2022; European Commission, 2017). To address these challenges effectively, the EU must prioritize actions for threatened species based on urgency, national responsibility, and the potential for synergies with broader environmental and climate goals (European Commission, 2021). Such an approach would also align with the Kunming-Montreal Global Biodiversity Framework, which aims to reverse ecosystem degradation through synergistic, nature-based solutions (McGowan et al. 2024).

A key component of the EU Biodiversity Strategy for 2030 is to ensure that species of conservation concern achieve or maintain a favourable conservation status (FCS) (EUR-Lex, 2020). In EU legislation, conservation status integrates both ecological and legal aspects to assess whether a species has sufficient habitat quality and quantity to sustain it as a long-term, functional part of the landscape (Oswald et al. 2025). Species of conservation concern are listed under the Habitats Directive (European Union Council, 1992), categorized into local populations within EU Member States jurisdictions and biogeographical regions (hereafter MS-BIOs; Figure 1). According to Article 17 of the Habitats Directive, Member States must review conservation status every six years (European Environment Agency, 2025). Ensuring comparability of conservation statuses across species is essential for identifying which local populations are effectively in FCS and which are not. This facilitates the identification of priority areas for additional protection, in support of conservation status improvement commitments (European Commission, 2021), while also preventing the misallocation of resources, such as disproportionately allocating funding to the protection of charismatic species (Mammola et al. 2020). However, the lack of clear methodological guidance for estimating FCS—apart from a few well-monitored species groups (Linnell & Boitani 2025)—forces EU Member States to rely on subjective criteria and expert judgment, which are often inconsistent and difficult to replicate, thereby undermining the framework’s effectiveness (Bijlsma et al. 2019).

**Figure 1.**
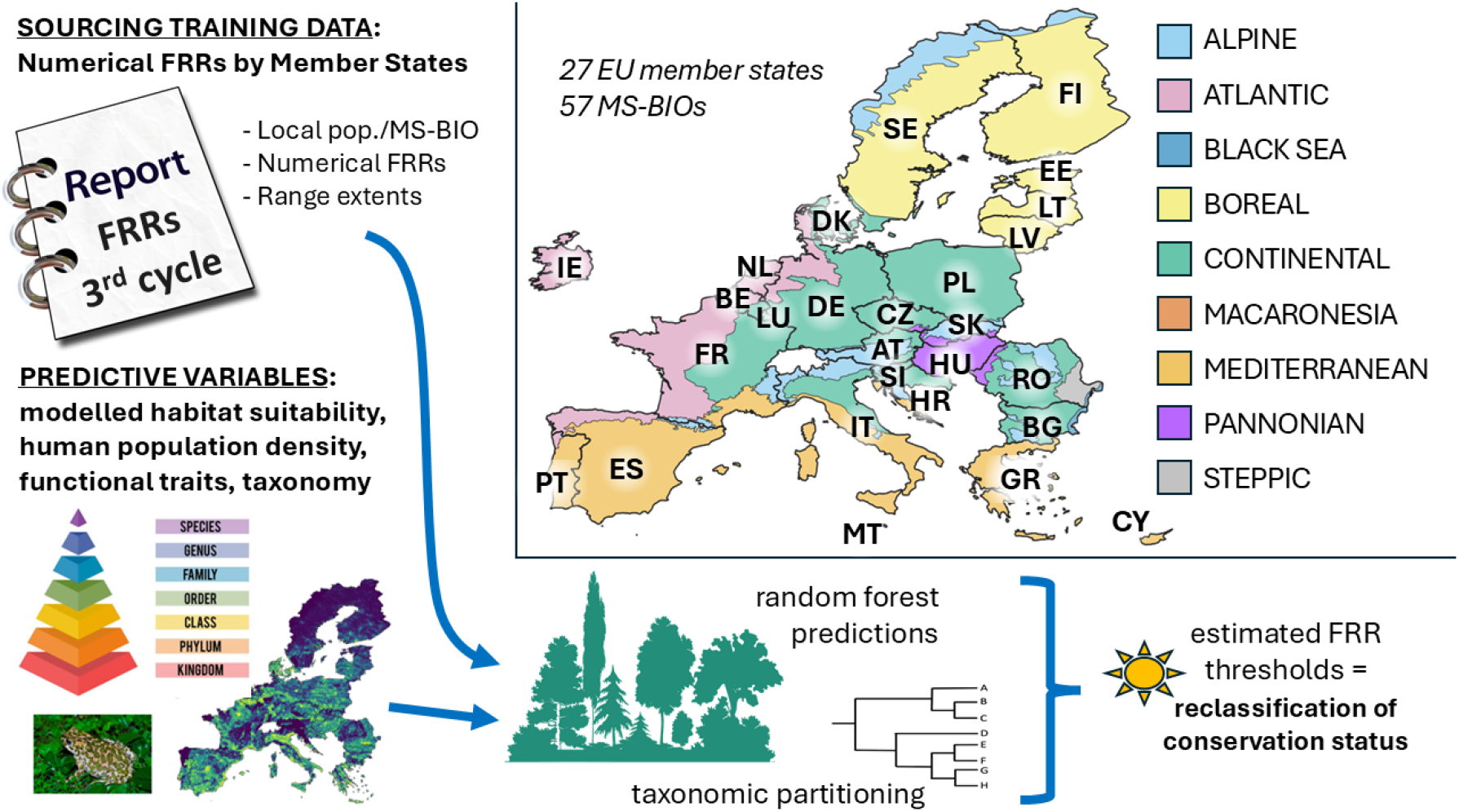
Top-right: geography of the 57 MS-BIO areas across 27 EU Member States (Table 2). Left and bottom: summary of the methods. N.B. Macaronesia is not included in the map. This biogeographical region covers the Spanish and Portuguese islands in the Atlantic Ocean.

A key metric for assessing the conservation status of local populations is the Favorable Reference Range (FRR), which defines the minimum range extent (km^2^) required to achieve FCS (Bijlsma et al. 2019). Estimating FRR is species-specific and relies on population monitoring, expert knowledge, historical baselines, and modelling techniques (Bonelli et al. 2021; D’Alessio et al. 2025). The criteria outlined in the Habitats Directive thus far have allowed using qualitative indicators instead of numerical FRR, including in the most recent 3^rd^ reporting cycle (period 2013-2018). Unlike a numerical FRR, a qualitative indicator assesses whether a population’s range extent is sufficient for FCS classification without defining a quantitative target. However, this approach limits precise conservation planning and hinders cross-species and MS-BIO comparisons (Bijlsma et al. 2019). In the 3^rd^ reporting cycle, most Member States relied heavily on qualitative indicators, with numerical FRRs missing for most local populations. This trend is largely due to insufficient data, often making expert opinion the primary basis for assigning conservation status (van Eldik et al. 2024). Addressing gaps in numerical FRRs is both urgent and essential, as these data offer far greater utility than qualitative indicators. Furthermore, with qualitative indicators set to be phased out in the upcoming 4^th^ reporting cycle (European Environment Agency, 2025), developing a robust method to establish numerical FRRs is a pressing priority.

To address this need, we developed a method that uses the limited numerical FRRs provided by Member States to estimate FRRs for as many local populations as possible. The approach employs machine learning to create cross-species comparisons based on habitat quality, functional traits, and taxonomy, filling gaps in reported FRRs and standardizing them for better comparability. We used the numerical FRRs reported for the 3^rd^ reporting cycle as our primary data source. Applying this methodology, we reassessed the conservation status of wildlife in the EU, offering a more accurate representation of biodiversity under this key metric in EU legal assets for nature protection. Our analysis focused on amphibians, mammals, and reptiles, but the methodology can be adapted to other taxa if similar data are available.

## 2 Materials and Methods

We collated data from the 3^rd^ reporting cycle (period 2013-2018) mandated by Article 17 of the Habitats Directive, linking taxonomy to each local population (Section 2.1). Next, we generated a set of predictive variables (Section 2.2), which were combined with numerical FRRs provided by Member States to estimate FRRs for as many local populations as possible within the relevant taxa, employing machine learning and taxonomic partitioning techniques (Section 2.3). Importantly, the developed modelling framework is not designed to predict species-specific trends in the response variable, i.e., FRR; instead, it is used to make FRR predictions by comparing local populations, akin to a gap-filling approach. Using the estimated FRRs, we reclassified the conservation status of local populations, assigning FCS if their range extent met or exceeded the FRR, and classifying them as unfavorable otherwise (Figure 1). Additionally, we calculated the proportional distance (p.dist) between each local population’s range extent and its estimated FRR, a mandatory indication from the upcoming 4^th^ reporting cycle (European Environment Agency, 2025). All analyses were performed in R v4.4.1 (R Core Team, 2024).

### 2.1 Gathering and processing FRRs reported by Member States

We downloaded the latest published version (3^rd^ reporting cycle, period 2013-2018) of the dataset on conservation status of habitat types and species under the Article 17 of the Habitats Directive (European Environment Agency datahub; Table S1). Local populations from marine biogeographical regions were excluded to preserve the consistency of the analyses conducted on the selected taxa, grouping predominantly terrestrial and freshwater species. Taxonomic classification (levels: class, order, family) was assigned by matching species names with the taxonomic backbone of the Global Biodiversity Information Facility (GBIF; Lane & Edwards 2007) using the R package taxize (Chamberlain et al. 2017b), without any manual modification. We retained only local populations of amphibians (class: *Amphibia*), mammals (class: *Mammalia*), and reptiles (orders: *Squamata* and *Testudines*), selecting 2,943 out of 8,097 total local populations. The excluded local populations belong to other taxa included in the same dataset, e.g., vascular plants. To ensure consistency, we constrained ‘range extent’ values and numerical FRRs to the spatial extent of the relevant MS-BIO, referencing a 1 km^2^ resolution raster map (ETRS89-extended/LAEA Europe equal-area projection). Furthermore, we standardized ‘range extent’ values and numerical FRRs as a percentage of MS-BIO area occupied, to address the variability in MS-BIO area extent that influences potential range extent and FRR. Finally, we extracted numerical FRRs assigned by Member States, which were the training data for our FRR estimates.

### 2.2 Predictive variables

The generated predictive variables fall into two groups: those related to A) habitat characteristics, and B) functional traits (Table 1). Importantly, not all predictive variables were available for every local population (Table S1).

**Table 1.** Predictive variables used in the Random Forest models to estimate FRRs. For both habitat characteristics and functional traits variables, the ecological hypothesis is that local populations with similar functional and life-history traits, occurring in comparable environments and disturbance regimes, are expected to have similar range extent needs to secure sufficient resources for long-term viability.

| CATEGORY | NAME | DATA SOURCE AND ELABORATION |
| --- | --- | --- |
| <u>Habitat characteristics</u> | median climate suitability | GBIF occurrence records combined with climate data (cc. 1 km resolution; annual mean 1979-2018; source: Copernicus Climate Change Service) reduced via Principal Component Analysis. Median value extracted across the relevant MS-BIO from habitat suitability models (Appendix 1). |
|  | climate suitability inequality | Gini index of climate suitability values modelled across the relevant MS-BIO. |
|  | median land use suitability | GBIF occurrence records combined with land cover data (100 m resolution; 2018 figure; source: Copernicus Land Monitoring Service) transformed to continuous scale. Median value extracted across the relevant MS-BIO from habitat suitability models (Appendix 1). |
|  | land use suitability inequality | Gini index of land-use suitability values modelled across the relevant MS-BIO. |
|  | available suitable habitat | Percentage of 1,000 randomly extracted 1 km<sup>2</sup> cells classified as suitable in the binary climate and land-use suitability models within the relevant MS-BIO. |
|  | median human population density | GBIF occurrence records interpolated with population density data map (5 km resolution; 2020 figure; source: Copernicus Emergency Management Service). Median value extracted across the relevant MS-BIO. |
|  | human population density inequality | Gini index of population density values at occurrence record points within the MS-BIO. |
| <u>Functional traits</u> | body mass | Measured or estimated (via trait-based correlation) in grams (log10-scaled for mammals). |
|  | litter size | Measured or estimated (via trait-based correlation) as the number of individuals (eggs for amphibians and reptiles) per reproductive event. |
|  | generation length | Measured or estimated (using taxonomy) as the average age of parents of the current cohort, in days. |

**Table 2.** Estimated FRRs and mean Nagelkerke’s R^2^ from RF regressions for each MS-BIO (all taxa included). Numbers in brackets indicate numerical FRRs provided by Member States relative to the total number of local populations per MS-BIO.

| MS-BIO | FRRs | $R^2$ | MS-BIO | FRRs | $R^2$ | MS-BIO | FRRs | $R^2$ |
| --- | --- | --- | --- | --- | --- | --- | --- | --- |
| AT-ALP (0/62) | 62 | 0.78 | ES-MED (4/85) | 85 | 0.81 | MT-MED (0/16) | 16 | 0.72 |
| AT-CON (1/62) | 62 | 0.78 | FI-ALP (0/9) | 9 | 0.68 | NL-ATL (0/43) | 42 | 0.74 |
| BE-ATL (0/39) | 38 | 0.72 | FI-BOR (0/30) | 29 | 0.72 | PL-ALP (0/47) | 46 | 0.69 |
| BE-CON (0/38) | 37 | 0.70 | FR-ALP (0/73) | 73 | 0.77 | PL-CON (2/52) | 51 | 0.74 |
| BG-ALP (69/70) | 70 | 0.77 | FR-ATL (0/64) | 64 | 0.81 | PT-ATL (20/46) | 46 | 0.72 |
| BG-BLS (66/72) | 72 | 0.75 | FR-CON (1/64) | 64 | 0.79 | PT-MAC (2/6) | 5 | 0.67 |
| BG-CON (85/90) | 90 | 0.77 | FR-MED (0/81) | 81 | 0.79 | PT-MED (22/56) | 56 | 0.79 |
| CY-MED (0/32) | 31 | 0.71 | GR-MED (0/112) | 112 | 0.78 | RO-ALP (0/56) | 56 | 0.70 |
| CZ-CON (0/61) | 61 | 0.75 | HR-ALP (0/63) | 63 | 0.72 | RO-BLS (0/20) | 20 | 0.87 |
| CZ-PAN (2/52) | 52 | 0.69 | HR-CON (0/65) | 65 | 0.73 | RO-CON (0/74) | 74 | 0.74 |
| DE-ALP (12/40) | 40 | 0.72 | HR-MED (0/76) | 76 | 0.75 | RO-PAN (0/31) | 31 | 0.78 |
| DE-ATL (14/45) | 44 | 0.81 | HU-PAN (0/67) | 67 | 0.76 | RO-STE (0/58) | 58 | 0.75 |
| DE-CON (28/58) | 58 | 0.82 | IE-ATL (13/15) | 15 | 0.76 | SE-ALP (11/12) | 12 | 0.74 |
| DK-ATL (0/22) | 22 | 0.79 | IT-ALP (0/73) | 73 | 0.75 | SE-BOR (34/34) | 33 | 0.78 |
| DK-CON (0/32) | 32 | 0.81 | IT-CON (0/69) | 69 | 0.76 | SE-CON (36/39) | 38 | 0.80 |
| EE-BOR (1/32) | 32 | 0.71 | IT-MED (0/94) | 94 | 0.78 | SI-ALP (25/64) | 64 | 0.73 |
| ES-ALP (1/54) | 54 | 0.72 | LT-BOR (0/37) | 36 | 0.71 | SI-CON (20/71) | 71 | 0.75 |
| ES-ATL (2/60) | 60 | 0.80 | LU-CON (12/38) | 37 | 0.71 | SK-ALP (0/60) | 60 | 0.71 |
| ES-MAC (16/23) | 23 | 0.67 | LV-BOR (22/40) | 39 | 0.68 | SK-PAN (0/59) | 59 | 0.72 |

We generated habitat characteristic variables by sourcing species occurrences (temporal range 2013–2018, corresponding to the 3^rd^ reporting cycle) from GBIF (DOI: https://doi.org/10.15468/dl.gxfms8; accessed 22/03/2025), collated and processed via the *R package rgbif* (Chamberlain et al. 2017a). These data were then used to determine the ‘median human population density’ where the local population occurs based on the occurrence data within the MS-BIO, and to run independent climate and land-use suitability models using an ensemble of machine learning and tree-based algorithms via the *R package sdm* (Naimi & Araújo 2016) (Appendix 1 and Appendix 2; ODMAP Protocol format, Zurell et al. 2020). The suitability models provided estimates for ‘median climate suitability’ and ‘median land-use suitability’ across the relevant MS-BIO, with the median extracted after predicting habitat suitability at all points within the MS-BIO, omitting spatial projection to minimize computational load. For human population density and the two suitability indices, we also calculated ‘distribution inequality’ using the Gini Index (Farris 2010), which measures the degree of disparity in the distribution of suitability values across an MS-BIO. This was done based on the hypothesis that these additional comparative variables could better reflect shared environmental conditions of local populations across MS-BIOs. We combined the suitability results to calculate the ‘percentage of suitable areas’ within each MS-BIO by converting the climate-based model and the land-use-based model into binary form, using the 90^th^ percentile prevalence from the model-building data as the threshold for suitability (Liu et al. 2005), and identifying intersecting suitable climatic and land-use areas. We sampled 1,000 random locations within the MS-BIO to calculate the percentage of such locations within the relevant MS-BIO. In this case, we applied a prevalence-based method to reduce computational load, as previously noted in justification for omitting spatial projection of the habitat suitability models.

For the functional trait variables, we used the databases ‘AmphiBIO’ (Oliveira et al. 2017) for amphibians, ‘COMBINE’ (Soria et al. 2021) for mammals, ‘ReptTraits’ (Oskyrko et al. 2024) for reptiles, and Mancini et al. (2025) for both amphibians and reptiles. For all three taxa, the selected traits were body mass (in grams, log10 scaled for mammals), litter size (mean number of offspring per litter for mammals or number of eggs per clutch), and generation length (the average age of parents in the current cohort; Pacifici et al. 2013). These traits were chosen for their availability in the datasets, their significance in demographic processes, and their relevance in extinction risk analyses (Foden et al. 2019).

### 2.3 Estimating FRRs and FCS reclassification

We developed a series of Random Forest (RF) regression models followed by taxonomic partitioning (Figure 1), to predict FRR for as many local populations as possible, including FRR already assigned by Member States. RF models were built separately for each taxon, as functional traits are not comparable across taxa, with numerical FRRs provided by Member States as the response variable. Predictive variables included habitat characteristics and functional traits (Table 1). The available data for the variables varied across local populations, depending on occurrence record availability, the success of habitat suitability models (Appendix 1), and functional trait data availability. To account for potential variability in the training data, stemming from different Member States assigning FRRs without explicit methodological coordination, we added ‘member state’ as an extra variable in each model. Rather than treating it as a categorical factor (which risked overfitting due to few levels for some Member States), we transformed it into a numerical variable by calculating each member state’s effect size, using Cohen’s d metric with Hedges’ correction (Cohen 2013). Effect size quantifies the magnitude of the difference between groups by comparing their means to the expected value, providing a standardized measure of the difference.

In the multiple RF model runs, we progressively reduced predictors from the complete set to a minimum of five, thus allowing us to derive FRR estimates also with incomplete data. Before running each model, we checked for variable collinearity and excluded variables with a variance inflation factor > 5 (Zuur et al. 2010). Each iteration prioritized models based on performance, measured by Nagelkerke R^2^ or, when performance was equal, by the number of predictors used, thus favouring a higher number of metrics for cross-species comparison. RF models were trained using the *R package RandomForest* (Breiman et al. 2018), with hyperparameter tuning of ‘ntree’ (number of decision trees; tested range 100 to 500, increment by 100), ‘mtry’ (number of features considered for each split; 2 to number of predictors - 1, by 1), and ‘nodesize’ (minimum number of observations in a terminal node; tested 1, 5, and 10), to optimize out-of-bag error. Given the limited training data per model, parameters were kept within recommended ranges to prevent overfitting (Probst et al. 2019). To account for each model’s predictive uncertainty, we calculated the mean squared prediction error. We adjusted each prediction by adding or subtracting this value (Lu & Hardin 2021) providing a prediction interval in addition to the specific FRR estimate.

For cases where the RF models did not provide estimates, we applied taxonomic partitioning following Pacifici et al. (2013) assigning remaining FRRs based on the mean among already-estimated FRRs within the relevant taxonomic group in the MS-BIO. Priority was given to estimates from populations within the same family, followed by order and class. The estimation error in this case was calculated using the standard deviation from the mean.

Local populations with an estimated FRR lower than or equal to their range extent were classified in FCS. In contrast, others were classified in unfavorable conservation status or labeled ‘N/D’ (Not Determined) if FRR estimation was impossible. Estimated FRRs were converted from the percentage of MS-BIO occupied area to km^2^ and capped to the extent of the MS-BIO of relevance. Finally, in accordance with the provisions of the 4^th^ reporting cycle (European Environment Agency, 2025), we calculated the proportional deviation between the estimated range and the FRR for all local populations classified in unfavourable conservation status.

## 3 Results

### 3.1 Estimated FRRs

We estimated a total of 2,929 (99.5%) FRRs (Table S2), compared to the 521 (17.7%) numerical FRRs available in the 3^rd^ reporting cycle (Table S1). Our predictions covered nearly all amphibians (579/580), all mammals (1847/1847), and most reptiles (503/516). The predictive performance of the RF regressions was good to optimal, with an overall mean Nagelkerke’s R^2^ of 0.754 (maximum 0.872 in Romania-Black Sea, minimum 0.667 in Spain-Macaronesia; Table 2). RF regressions performed particularly well for amphibians (0.905) and reptiles (0.924), while results for mammals (0.669) were relatively less satisfactory. The majority of numerical FRRs were derived from RF regressions (82.7%), followed by family-level partitioning (7.5%), order-level partitioning (6.1%), and class-level partitioning (3.2%) (Figure 2).

**Figure 2.**
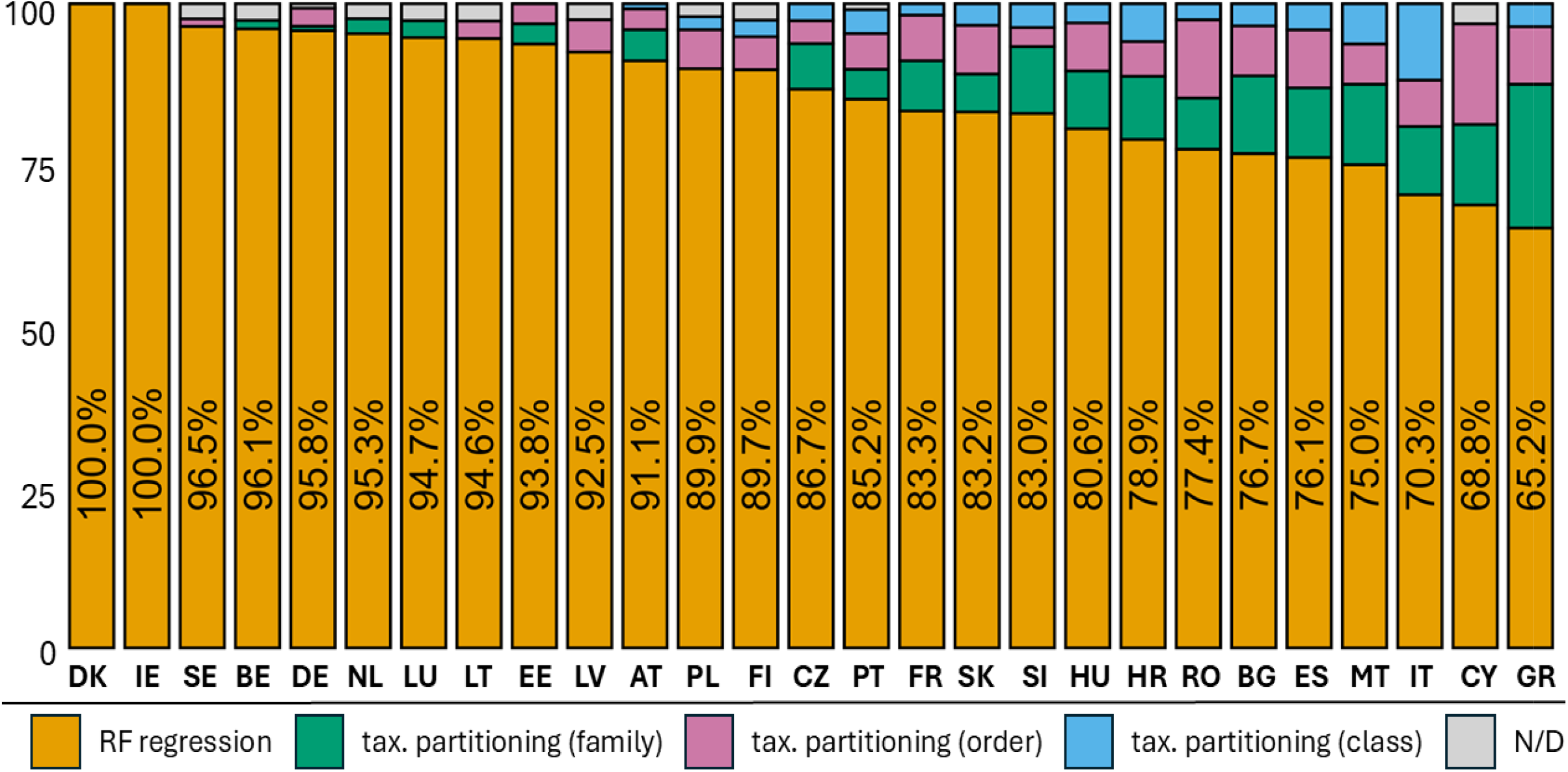
Percentage of estimated FRRs over the total number of local populations per Member State by methodology. Member States are ordered left to right by decreasing contribution of the RF regressions (percentage reported on the bar), which is the primary modelling approach.

### 3.2 Reclassification of conservation status and mean proportional distance to FCS

Based on our estimates, 1,024 of 2,943 local populations (34.8%) are in FCS. Considering RF prediction errors and taxonomic partitioning variability, this ranges from 939 (31.9%) to 1,099 (37.3%) (Table S2). In contrast, the Report indicates 69.1% in FCS considering both numerical FRRs and qualitative indicators. Discrepancies are most notable in Fennoscandia, Italy, and Eastern Europe, especially for amphibians and reptiles (Figure 3). Among populations in unfavourable status, reptiles show the largest mean proportional deviation between current range and their FRRs (-67.8%), followed by amphibians (-64.2%) and mammals (-63.4). A high share of local populations exceeds a -75% deviation, especially among amphibians and mammals. France, Italy, and Romania have the highest numbers of populations in unfavourable conservation status (Table 3).

**Table 3.** Number of local populations in unfavourable conservation status by Member State (totals in brackets) and proportional deviation (p.dev) between current range and estimated FRR, grouped by taxon—amphibians (Amp.), mammals (Mam.), reptiles (Rep.)—and organized by p.dev intervals. Colours indicate the p.dev interval with the highest number of unfavourable populations per taxon and Member State: green = amphibians, orange = mammals, blue = reptiles

| EU Member States | p.dev. < -0.25 |  |  | p.dev. -0.25 to -0.50 |  |  | p.dev. -0.50 to -0.75 |  |  | p.dev. -0.75 to -1.00 |  |  |
| --- | --- | --- | --- | --- | --- | --- | --- | --- | --- | --- | --- | --- |
|  | Amp. | Mam. | Rep. | Amp. | Mam. | Rep. | Amp. | Mam. | Rep. | Amp. | Mam. | Rep. |
| Austria (110) | 1 | 6 | 1 | 2 | 12 | 5 | 6 | 15 | 10 | 14 | 38 | 0 |
| Belgium (29) | 1 | 3 | 0 | 2 | 1 | 0 | 1 | 5 | 1 | 3 | 12 | 0 |
| Bulgaria (117) | 5 | 17 | 11 | 4 | 16 | 7 | 3 | 14 | 13 | 11 | 10 | 6 |
| Croatia (116) | 4 | 12 | 2 | 3 | 10 | 2 | 1 | 20 | 13 | 5 | 40 | 4 |
| Cyprus (9) | / | 1 | 1 | / | 1 | 0 | / | 1 | 2 | / | 2 | 1 |
| Czech Rep. (43) | 2 | 4 | 0 | 1 | 3 | 1 | 3 | 4 | 4 | 3 | 18 | 0 |
| Denmark (38) | 3 | 3 | 2 | 4 | 3 | 0 | 4 | 4 | 0 | 1 | 14 | 0 |
| Estonia (16) | 3 | 3 | 0 | 6 | 0 | 0 | 2 | 1 | 1 | 13 | 7 | 0 |
| Finland (18) | 1 | 1 | / | 0 | 1 | / | 0 | 7 | / | 1 | 7 | / |
| France (247) | 5 | 22 | 4 | 7 | 34 | 6 | 9 | 36 | 16 | 34 | 71 | 3 |
| Germany (91) | 9 | 17 | 1 | 4 | 10 | 0 | 5 | 9 | 8 | 5 | 22 | 1 |
| Greece (84) | 1 | 5 | 2 | 1 | 6 | 7 | 4 | 4 | 23 | 9 | 10 | 12 |
| Hungary (46) | 0 | 7 | 1 | 1 | 5 | 2 | 0 | 8 | 4 | 5 | 11 | 2 |
| Ireland (5) | 0 | 1 | / | 0 | 1 | / | 0 | 1 | / | 1 | 1 | / |
| Italy (182) | 5 | 11 | 1 | 7 | 21 | 4 | 7 | 14 | 28 | 27 | 50 | 7 |
| Latvia (7) | 0 | 0 | 0 | 0 | 1 | 0 | 2 | 0 | 1 | 5 | 5 | 1 |
| Lithuania (7) | 2 | 0 | 0 | 0 | 0 | 0 | 1 | 0 | 1 | 2 | 1 | 0 |
| Luxembourg (10) | 0 | 0 | / | 0 | 0 | / | 3 | 0 | / | 0 | 7 | / |
| Malta (10) | / | 0 | 0 | / | 1 | 0 | / | 3 | 1 | / | 4 | 1 |
| Netherlands (23) | 3 | 1 | 0 | 1 | 3 | 1 | 0 | 3 | 1 | 4 | 6 | 0 |
| Poland (66) | 3 | 2 | 0 | 3 | 8 | 1 | 1 | 14 | 1 | 4 | 29 | 0 |
| Portugal (60) | 1 | 16 | 2 | 0 | 8 | 0 | 1 | 15 | 4 | 1 | 10 | 2 |
| Romania (172) | 3 | 7 | 8 | 3 | 12 | 7 | 2 | 21 | 16 | 5 | 77 | 11 |
| Slovakia (116) | 1 | 6 | 0 | 7 | 18 | 7 | 4 | 17 | 6 | 10 | 39 | 1 |
| Slovenia (51) | 2 | 7 | 0 | 3 | 12 | 2 | 2 | 7 | 7 | 5 | 2 | 2 |
| Spain (40) | 3 | 5 | 4 | 6 | 19 | 8 | 2 | 12 | 26 | 13 | 33 | 8 |
| Sweden (139) | 2 | 7 | 0 | 3 | 12 | 2 | 0 | 7 | 7 | 4 | 2 | 2 |

**Figure 3.**
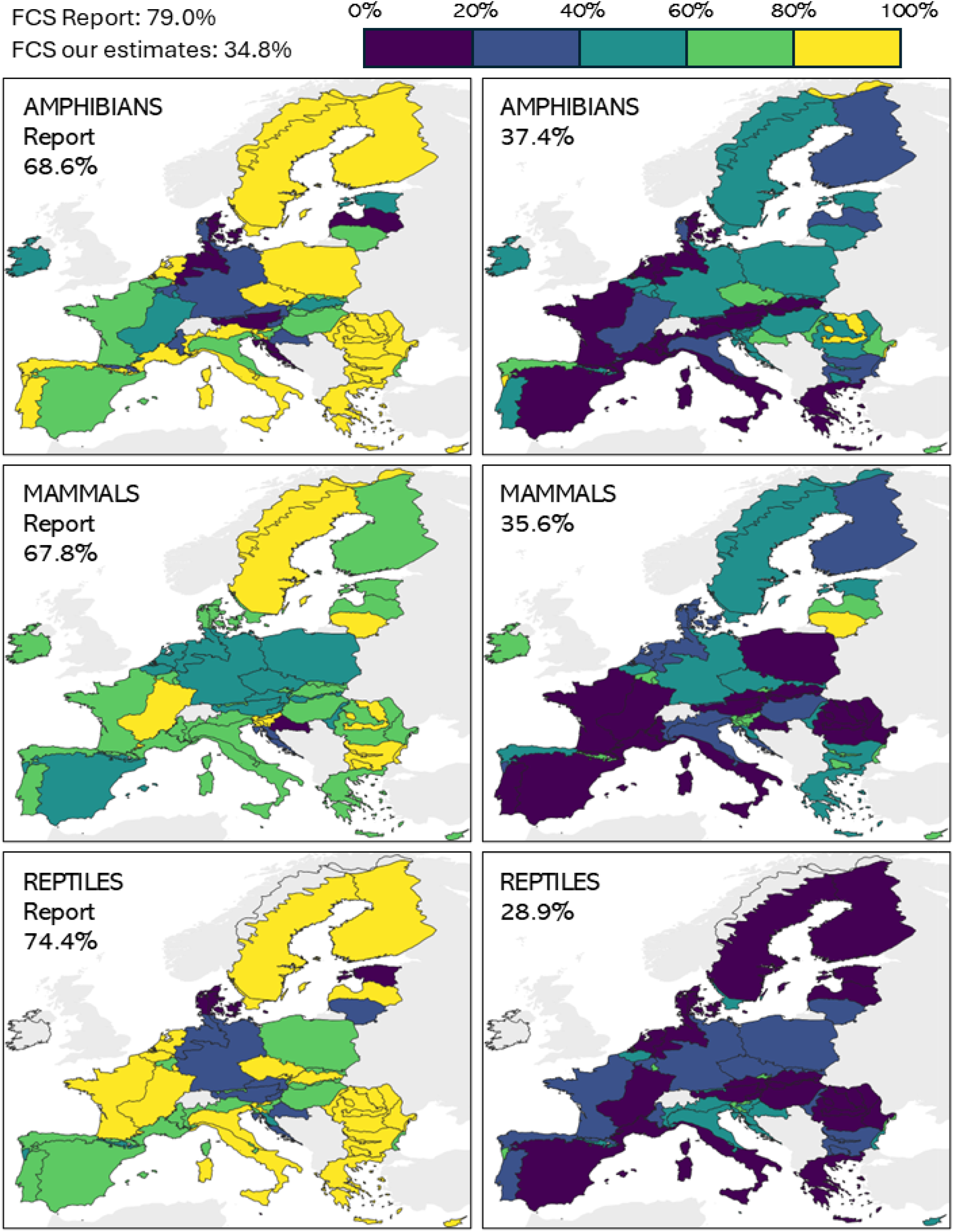
Left: proportion of local populations in FCS from the FRRs reported by Member States (values in brackets: percentage of local populations in FCS over the total). Right: proportion of local populations in FCS based on our reclassifications.

## 4 Discussion

Despite several conceptual frameworks proposed to improve coordination among EU Member States in the definition of FRR targets (Bijlsma et al. 2019; van Eldik et al. 2024), provided numerical FRRs remain scarce and likely inconsistent, as also noted in previous reports (Hochkirch et al. 2013). This is largely because no alternative methodologies departing from the prevailing paradigm have been adopted beyond species-specific approaches that require extensive ecological expertise and long-term monitoring data. In this study, we presented a novel framework based on cross-species comparisons of traits, habitat structure, and taxonomic partitioning, substantially expanding the availability of FRRs for species of conservation concern in the EU. Compared to the 3^rd^ reporting cycle, referencing the period 2013-2018, our method covers FRR for over 80% more local populations. At its core are machine-learning-based regression models, which achieved strong predictive performance, particularly in countries like Germany and Sweden, where species-specific numerical FRRs were available to train the models.

With FRRs now assigned to nearly all local populations, our findings diverge significantly from previous assessments of wildlife conservation status in the EU. While FRRs provided by Member States— based largely on qualitative indicators—estimated 69.1% of local populations in FCS, our analysis places this figure at just 34.8%. When accounting for model uncertainty, the estimate ranges from 31.9% to 37.3%, underscoring the robustness of our results. These findings are consistent with the growing evidence of widespread biodiversity decline across Europe, with only limited exceptions among mammalian and avian species (Burns et al. 2021; Warren et al. 2021; Ledger et al. 2022). Bijlsma et al. (2019) reported a similar discrepancy with reference to the 2^nd^ reporting cycle (2012–2017), noting that, considering qualitative indicators, approximately 60% of local populations (across all taxa of species of conservation concern) were considered to be in FCS. In contrast, when only numerical FRRs were considered, the trend indicated that for most local populations, the range extent was smaller than the FRR. Amphibians and mammals represent a large share of populations in unfavourable conservation status; however, reptiles show the greatest deviation from their FRRs (-67.8% on average). More broadly, across all three taxa, many populations not currently reaching FCS are significantly below their estimated FRR. This suggests that numerous local populations, likely endemic to specific regions, may have been severely affected by long-standing habitat fragmentation—particularly in Southern and Central Europe, where such processes have occurred for centuries (Kaplan et al. 2009). For these populations, protection of current habitats alone may be insufficient to achieve FCS. Active habitat restoration, combined with long-term habitat protection, may be necessary (Araújo & Alagador 2024). Alternatively, it may be necessary to re-evaluate the relevance of the FRR framework for such populations with highly localized ranges, for which conventional conservation targets may not reflect ecological realities.

Importantly, our approach is not intended to replace species-specific expert assessments but to complement them. Its main limitation lies in the need for a sufficiently large number of numerical FRRs already provided by Member States to serve as training data. As such, the method’s efficiency and reliability would greatly benefit from systematic coordination with national experts, who should focus on providing the most ecologically robust numerical FRR values possible. These experts could then apply our framework to estimate FRRs for species where ecological knowledge or long-term monitoring remains limited. As demonstrated, even with a starting dataset covering only 17% of species of conservation concern, the method can predict FRRs for almost all species, with strong model performance. Additional limitations include gaps in species occurrence data and missing functional ecological information, which can make our modelling unfeasible or unreliable. Another constraint arises from harmonising FRRs and range extents at the MS-BIO level, which risks assuming uniform ecological requirements across diverse regions. Although this was necessary to secure adequate training data, it may produce less reliable estimates for species highly dependent on region-specific habitat conditions. Future refinements could recalibrate FCS classifications at the EU level, integrating finer-scale ecological data where available. While alternative approaches —such as historical range reconstructions (Pacifici et al. 2019), climate-resilient baselines (Willis et al. 2010), or abundance-based models (D’Alessio et al. 2025)— hold conceptual value, they require data unavailable for most EU species of conservation concern.

We believe this work makes a meaningful contribution to advancing conservation practices in the EU by providing a fundamental tool that complements species-specific estimates of FRRs. Critically, it offers robust numerical FRR estimates, helping to reduce the longstanding overreliance on qualitative assessments —widely acknowledged as imprecise and recently excluded from EU reporting guidelines (European Environment Agency, 2025). The method also addresses persistent inconsistencies stemming from divergent interpretations of the FCS concept, including its occasional mischaracterisation as the minimum viable population (Epstein et al. 2016; Trouwborst et al. 2017) or disproportionate influence from species charisma (Brambilla et al. 2013). A key innovation of our framework is the integration of quantified model uncertainty alongside FRR estimates, offering pragmatic flexibility. This supports our view that FRRs should be seen not as fixed figures, but as policy-relevant broad indications or range extents to be pursued to ensure species of conservation concern have access to adequate habitat and resources.

