## Appendix 1 for "Improving the classification of wildlife conservation status to support nature protection in the European Union"

– ODMAP Protocol –

2025-05-21

### Overview

##### Authorship

<Contact >

<Study link>

##### Model objective

Model objective: Forecast and transfer

Target output: continuous habitat suitability index, binary habitat suitability index

##### Focal Taxon

Focal Taxon: Amphibians, mammals and reptiles of conservation concern under the European Union Habitats Directive (EU Council Directive 92/43/EEC; Annexes II and IV)

##### Location

Location: European Union (continent + Macaronesia)

##### Scale of Analysis

<Spatial extent>

Spatial resolution: 1

Temporal extent: 2013-2018

Boundary: political

##### Biodiversity data

Observation type: occurrence records (GBIF), human observations (GBIF), machine observations (GBIF)

Response data type: point occurrence

##### Predictors

Predictor types: climatic

##### Hypotheses

Hypotheses: Broad-scale climatic gradients primarily determine suitable habitat for a species.

##### Assumptions

Model assumptions: Biotic interactions not accounted for.

##### Algorithms

Modelling techniques: brt, maxlike, maxnet, ranger, rpart, svm

Model complexity: Machine learning and tree-based algorithms to model non-linear and complex relationships and to maximize predictive accuracy.

Model averaging: Mean of the predictions from models with an Area Under the Curve (AUC) score ≥ 0.7.

##### Workflow

Model workflow: For each species of conservation concern of the taxa considered, we predicted habitat suitability based on georeferenced occurrence records collected at a global scale. This broad-scale approach was adopted to capture the full potential environmental variability of each species and to minimize the risk of niche truncation (Chevalier et al., 2022). Habitat suitability was modeled using climatic predictors downloaded from the Copernicus Climate Change Service (averaged annual means of the period 1979-2018), reduced using Principal Component Analysis (PCA) to limit spatial correlation, retaining only principal components with eigenvalues > 1 (De Marco & Nobrega, 2018). We checked for any remaining multicollinearity among the principal components by calculating the Variance Inflation Factor (VIF) among the predictor variables, using a highly conservative threshold of 3 (Zuur et al., 2010). Correlated variables exceeding this threshold were removed. This procedure was repeated for each model run—thus, for each species—by extracting the values of the predictor variables at the species’ occurrence geographical coordinates. We employed machine learning and tree-based methods available in the R package sdm (Araujo & Naimi, 2016). As background or pseudo-absence data, we randomly selected ten points for each occurrence record considered. We then created an ensemble of projections from these algorithms, including only outputs with validation metrics exceeding 0.7 AUC. Validation was performed by splitting the occurrence records into 70% training data and 30% test data. Projections were made using the same climatic variables used as predictors. Subsequently, for each local population, we extracted the median predicted habitat suitability within its corresponding MS-BIO region, and used these values as predictors to model FRRs (see methods in main text). Finally, we binarized the climatic model output for each species into potential presence/absence using the 90% prevalence threshold method (Liu et al., 2005).

Chevalier, M., Zarzo-Arias, A., Guélat, J., Mateo, R. G., & Guisan, A. (2022). Accounting for niche truncation to improve spatial and temporal predictions of species distributions. Frontiers in Ecology and Evolution, 10, 944116.

De Marco, P., & Nóbrega, C. C. (2018). Evaluating collinearity effects on species distribution models: An approach based on virtual species simulation. PloS one, 13(9), e0202403.

Zuur, A. F., Ieno, E. N., & Elphick, C. S. (2010). A protocol for data exploration to avoid common statistical problems. Methods in ecology and evolution, 1(1), 3-14.

Naimi, B., & Araújo, M. B. (2016). sdm: a reproducible and extensible R platform for species distribution modelling. Ecography, 39(4), 368-375.

Liu, C., Berry, P. M., Dawson, T. P., & Pearson, R. G. (2005). Selecting thresholds of occurrence in the prediction of species distributions. Ecography, 28(3), 385-393.

##### Software

Software: Platform: R v4.4.1 (R Core Team, 2024); Key packages: ‘enmSdmX’ (v1.2.12), ‘sdm’ (v1.2.55), ‘usdm’ (v2.1.7)

Code availability: <https://figshare.com/s/fff905545511bd5b4e78>

<Data availability>

### Data

##### Biodiversity data

Taxon names: amphibians, mammals, reptiles

Taxonomic reference system: the taxonomic backbone of GBIF using the R package taxize (v0.9.100)

Ecological level: species

Data sources: <https://doi.org/10.15468/dl.gxfms8>

Sampling design: opportunistic occurrence records, random background records

Sample size: Limited to 200 occurrence records, retaining only one per spatial resolution unit of the climatic data, randomly selected to reduce computation time. 10 background points for each occurrence record

Clipping: European Union

Cleaning: retained only occurrence records with coordinates uncertainty < 1000m (1km), from GBIF sources ‘occurrence records’, ‘human observations’, ‘machine observations’

<Absence data>

Background data: For each species, ten points were randomly selected for each occurrence record considered.

##### Data partitioning

Training data: 70% occurrence and background records

Validation data: 30% occurrence and background records

Test data: truly independent data , sensu Hastie, et al. (2009)

Hastie, T., Tibshirani, R., Friedman, J., Hastie, T., Tibshirani, R., & Friedman, J. (2009). Unsupervised learning. The elements of statistical learning: Data mining, inference, and prediction, 485-585. 30% occurrence and background records

##### Predictor variables

Predictor variables: Annual mean temperature (BIO01), Annual precipitation (BIO12), Aridity annual mean, Cloud cover, Evaporative fraction annual mean, Growing degree days, Isothermality (BIO03), Maximum temperature of the warmest month (BIO05), Mean diurnal range (BIO02), Mean temperature of coldest quarter (BIO11), Mean temperature of driest quarter (BIO09), Mean temperature of warmest quarter (BIO10), Mean temperature of wettest quarter (BIO08), Minimum temperature of the coldest month (BIO06), Potential evaporation annual mean, Precipitation of coldest quarter (BIO19), Precipitation of driest month (BIO14), Precipitation of driest quarter (BIO17), Precipitation of warmest quarter (BIO18), Precipitation of wettest month (BIO13), Precipitation of wettest quarter (BIO16), Precipitation seasonality (BIO15), Surface latent heat flux annual mean, Surface sensible heat flux annual mean, Temperature annual range (BIO07), Temperature seasonality (BIO04), Volumetric soil water layer 1 annual mean, Water vapour pressure

Data sources: Copernicus Climate Data Store DOI: 10.24381/cds.bce175f0

<Spatial extent>

Spatial resolution: 0.5° x 0.5°

Coordinate reference system: EPSG:4326 – WGS 84

<Temporal extent>

Data processing: PCA analysis to derive uncorrelated principal components with eigenvalues > 1

##### Transfer data

Data sources: <https://cds.climate.copernicus.eu/datasets/sis-biodiversity-era5-global?tab=overview> Version 1.0

<Spatial extent>

<Spatial resolution>

<Temporal extent>

<Models and scenarios>

<Quantification of Novelty>

### Model

##### Multicollinearity

Multicollinearity: Retained only uncorrelated variables (VIF > 3)

##### Model settings

<brt>

<maxlike>

<maxnet>

<ranger>

<rpart>

<svm>

<Model settings (extrapolation)>

##### Model estimates

Coefficients: median posterior

##### Analysis and Correction of non-independence

<Spatial autocorrelation>

##### Threshold selection

<Threshold selection>

### Assessment

##### Performance statistics

<Performance on training data>

<Performance on validation data>

<Performance on test data>

##### Plausibility check

<Response shapes>

<Expert judgement>

### Prediction

##### Prediction output

Prediction unit: Habitat Suitability Index

##### Uncertainty quantification

<Scenario uncertainty>

<Novel environments>
