## Appendix 2 for "Improving the classification of wildlife conservation status to support nature protection in the European Union"

– ODMAP Protocol –

2025-05-22

### Overview

##### Authorship

<Contact >

<Study link>

##### Model objective

Model objective: Forecast and transfer

Target output: continuous habitat suitability index, binary habitat suitability index

##### Focal Taxon

Focal Taxon: Amphibians, mammals and reptiles of conservation concern under the European Union Habitats Directive (EU Council Directive 92/43/EEC; Annexes II and IV)

##### Location

Location: European Union (continent + Macaronesia)

##### Scale of Analysis

<Spatial extent>

Spatial resolution: 1

Temporal extent: 2013-2018

Boundary: political

##### Biodiversity data

Observation type: occurrence records (GBIF), human observations (GBIF), machine observations (GBIF)

Response data type: point occurrence

##### Predictors

Predictor types: habitat

##### Hypotheses

Hypotheses: Species tolerance to land use and preferences for land cover (e.g., a particular forest type) determines habitat suitability at a broad scale.

##### Assumptions

Model assumptions: Biotic interactions not accounted for.

##### Algorithms

Modelling techniques: brt, maxlike, maxnet, ranger, rpart, svm

Model complexity: Machine learning and tree-based algorithms to model non-linear and complex relationships and to maximize predictive accuracy.

Model averaging: Mean of the predictions from models with an Area Under the Curve (AUC) score ≥ 0.7.

##### Workflow

Model workflow: For each species of conservation concern of the taxa considered, we predicted habitat suitability based on georeferenced occurrence records collected across Europe. This contrasts with the collection of records at a global scale done for the climatic habitat suitability model, as for land use, we were particularly interested in species’ tolerance levels specifically in relation to European Union socio-ecological dynamics. Habitat suitability was modeled using land-use predictors downloaded from the Copernicus Land Monitoring Service (CORINE land cover, 2018 figure). The categorical CORINE variables were converted to continuous scale while downscaling the resolution from the original 100 meters to the working resolution of 1 kilometer, by calculating the percentage of cells belonging to each land cover category within the new resolution grid, which were then used as predictor components. We checked for multicollinearity among the predictors by calculating the Variance Inflation Factor (VIF) among the predictor variables, using a highly conservative threshold of 3 (Zuur et al., 2010). Correlated variables exceeding this threshold were removed. This procedure was repeated for each model run—thus, for each species—by extracting the values of the predictor variables at the species’ occurrence geographical coordinates. We employed machine learning and tree-based methods available in the R package sdm (Araujo & Naimi, 2016). As background or pseudo-absence data, we randomly selected ten points for each occurrence record considered. We then created an ensemble of projections from these algorithms, including only outputs with validation metrics exceeding 0.7 AUC. Validation was performed by splitting the occurrence records into 70% training data and 30% test data. Projections were made using the same land use variables used as predictors. Subsequently, for each local population, we extracted the median predicted habitat suitability within its corresponding MS-BIO region, and used these values as predictors to model FRRs (see methods in main text). Finally, we binarized the climatic model output for each species into potential presence/absence using the 90% prevalence threshold method (Liu et al., 2005).

##### Predictor variables

Predictor variables: ‘Percentage of croplands’ (levels 12:17, 19:20 in original format of CORINE land cover categories), ‘percentage of pastures’ (level 18 in original format), ‘percentage of agri-forest areas’ (levels 21:22 in original format), ‘broad-leaved forests’ (level 23 in original format), ‘coniferous forests’ (level 24 in orginal format), ‘mixed-forests’ (level 25 in original format), ‘grasslands’ (level 26 in original format), ‘shrublands’ (levels 27:29 in original format), ‘wetlands’ (levels 35:43 in original format).

Data sources: Copernicus Land Monitoring Service, CORINE Land Cover 2018; DOI: <https://doi.org/10.2909/960998c1-1870-4e82-8051-6485205ebbac>

<Spatial extent>

Spatial resolution: 100m x 100m

Coordinate reference system: EPSG:3035

<Temporal extent>

Temporal resolution: 2017-2018

Data processing: Rasters were rescaled to a resolution of 1 km by transforming categorical variables into continuous ones, calculating, for each category, the percentage of original cells composing the area corresponding to the rescaled 1 km resolution.

##### Transfer data

<Data sources>

<Spatial extent>

<Spatial resolution>

<Temporal extent>

<Models and scenarios>

<Quantification of Novelty>

### Model

##### Multicollinearity

Multicollinearity: Retained only uncorrelated variables (VIF > 3)

##### Model settings

<brt>

<maxlike>

<maxnet>

<ranger>

<rpart>

<svm>

<Model settings (extrapolation)>

##### Model estimates

Coefficients: median posterior

##### Analysis and Correction of non-independence

<Spatial autocorrelation>

##### Threshold selection

<Threshold selection>

### Assessment

##### Performance statistics

<Performance on training data>

Performance on validation data: AUC

Performance on test data: AUC

##### Plausibility check

<Response shapes>

<Expert judgement>

### Prediction

##### Prediction output

Prediction unit: Habitat Suitability Index

##### Uncertainty quantification

<Scenario uncertainty>

<Novel environments>
