## Supplementary Text for "Improving the classification of wildlife conservation status to support nature protection in the European Union"

**SUPPLEMENTARY INFORMATION**

### **Climate-based habitat suitability model**

Per ogni specie di conservation concern dei taxa considerati abbiamo predetto habitat suitability basandoci su occurrence records raccolti a scala globale (per considerare tutto ik potenziale di variabilita ambientale della specie dunque limitare niche truncation) basandoci su predittori climatici: a seguire, considerando ogni popolazione locale, abbiamo estratto il valore mediano dalle predizioni di habitat suitability nella MS-BIO di appartenenza. We obtained georeferenced occurrence records per ogni specie di conservation concern from GBIF. Data were cleaned by keeping solo occurrence records con una incertezza di posizione pari alla risoluzione dello studio (1000m – 1km). Inoltre, abbiamo selezionato solo occurrence records del periodo 2013-2018, corrispondenti al periodo di riferimento per il 3rd ciclo del report habitats directive, usato come dati training per i nostri modelli di stima dei valori FRR. Inoltre, abbiamo tenuto solo occurrence records provenienti dalle categorie GBIF ‘human observation’, ‘occurrence’, ‘machine observation’. Abbiamo scaricato da Copernicus le seguenti variabili climatiche: XXX, XXX, XXX, and XXX. Tutte queste variabili sono state ridotte in variabili derivate tramite principal component analysis (PCA), mantenendo unicamente le variabili derivati con eigenvalues > 1 ()REF

**2. Occurrence Data**

Species occurrence records were obtained from *[source, e.g., GBIF, field surveys]* and included *[number]* georeferenced presence records. Data were cleaned to remove duplicates, spatial outliers, and records with missing coordinates. For presence-absence models, absence or pseudo-absence data were generated using *[method, e.g., random background sampling, target-group background]*.

**3. Environmental Variables**

Environmental predictors included *[list of variables, e.g., temperature, precipitation, elevation]* obtained from *[source, e.g., WorldClim v2, CHELSA]* at a spatial resolution of *[e.g., 1 km²]*. Variables were selected based on ecological relevance and reduced for multicollinearity using *[e.g., Variance Inflation Factor (VIF), Pearson correlation]*.

**4. Modeling Algorithm**

The species distribution was modeled using *[model type, e.g., MaxEnt, Random Forest, Generalized Linear Model]*, which is appropriate for *[data type: presence-only / presence-absence]* data. The model was trained on *[percentage]* of the data and evaluated using the remaining *[percentage]* as test data.

**5. Model Evaluation**

Model performance was assessed using *[evaluation metrics, e.g., AUC, TSS, Kappa]*. The final model achieved an AUC of *[value]*, indicating *[interpretation of performance]*. Variable importance was examined to identify the key environmental drivers of the species’ distribution.

**6. Projection and Mapping**

The model was projected across the study area to generate a habitat suitability map, highlighting areas of high and low likelihood of species presence. Suitability scores ranged from 0 (unsuitable) to 1 (highly suitable). Maps were visualized using *[software, e.g., QGIS, ArcGIS, R]*.

**7. Limitations and Assumptions**

The model assumes that the species is at equilibrium with its environment and that occurrence records represent true presence. Limitations include potential sampling bias, coarse spatial resolution of environmental data, and the lack of biotic interaction variables.
